# Corticocortical Connections of the Rostral Forelimb Area in Rats: A Quantitative Tract-Tracing Study

**DOI:** 10.1101/2020.06.10.144923

**Authors:** Edward T Urban, Mariko Nishibe, Scott Barbay, David J Guggenmos, Randolph J Nudo

## Abstract

The rostral forelimb area (RFA) in the rat is considered to be a premotor cortical region based primarily on its efferent projections to the primary motor cortex. The purpose of the present study was to identify corticocortical connections of RFA, and to describe the relative strength of connections with other cortical areas. This will allow us to better understand the broader cortical network in which RFA participates, and thus, determine its function in motor behavior. In the present study, the RFA of adult male Long-Evans rats (n=6) was identified using intracortical microstimulation techniques and injected with the tract tracer, biotinylated dextran amine (BDA). In post-mortem tissue, location of BDA-labeled terminal boutons and neuronal somata were plotted and superimposed on cortical field boundaries. The results demonstrated that the RFA has dense to moderate reciprocal connections with primary motor cortex, the frontal cortex medial and lateral to RFA, primary somatosensory cortex (S1), and lateral somatosensory areas. Importantly, S1 connections were dense to moderate in dysgranular zones, but sparse to negligible in granular zones. Cortical connections of RFA in rat are strikingly similar to cortical connections of the ventral premotor cortex in non-human primates, suggesting that these areas share similar functions.

---

The motor areas of the cerebral cortex of mammals generally have been divided into the primary motor cortex (M1) and various premotor areas. By definition, premotor areas provide direct input to M1 (Dum RP and PL Strick 2002). While several premotor areas have been identified in non-human primates, the status of premotor cortex in rodent species is still unclear. Rodents are used increasingly in studies focused on understanding the cortical control of movement, and the role of premotor areas in recovery after injury to M1 (Urban ET, 3rd et al. 2012; Nishibe M et al. 2015; Tennant KA et al. 2015; Touvykine B et al. 2015). Therefore, it is important to understand the relationship of premotor areas in rodents and primates, and the similarity of their anatomical connectivity with other cortical areas.

Functionally, M1 is thought to be in the most direct control of movements of skeletal musculature. While the non-primary motor areas also have descending connections to the spinal cord (Nudo RJ and RB Masterton 1990), premotor cortex is thought to be involved with higher order processing, affecting motor control primarily by modulating M1 (Dum RP and PL Strick 2002). Numerous interconnections between premotor areas and M1 could relay information about movement selection, preparation for movement, generation of movement sequences, or sensory guidance of movement. Thus, the cortical connectivity of premotor areas may provide critical information regarding the information flow within the cortex, and thus, the putative role that premotor areas play in sensorimotor control in rodents.

In rats, cortical motor areas have been defined on the basis of both cytoarchitecture and the motor response to electrical stimulation (Donoghue JP and SP Wise 1982). Using intracortical microstimulation (ICMS) techniques, two motor forelimb fields can be identified. The larger field is the region from which movements can be evoked by the lowest intensity of ICMS current. Cytoarchitechtonically, this zone is identified as the lateral agranular field containing large layer 5 neurons. Because of its similarity to M1 in primates, this area is typically considered to be the rat’s M1. The forelimb representation in rat M1 has been most studied, and is called the caudal forelimb area (CFA).

A second region from which forelimb movements can be evoked by ICMS has been called the rostral forelimb area (RFA). On the basis of ICMS results, this area is separated from CFA by a thin strip of cortex where ICMS elicits neck or whisker movements. While cytoarchitectonically distinct from the CFA, layer 5 neurons in both areas send fibers to the spinal cord which terminate in the intermediate zone and ventral horn (Rouiller EM et al. 1993), much like frontal motor areas in non-human primates. Similar to M1 and premotor cortex of non-human primates, CFA and RFA are strongly and reciprocally interconnected by corticocortical fibers.

These anatomical and physiological commonalities have led several investigators to propose that the RFA is either the rat premotor cortex, or the supplementary motor area (Rouiller EM *et al*. 1993; Nudo RJ and SB Frost 2007). One approach to compare the functional importance of cortical areas in different species is to understand similarities and differences in the connection patterns with other cortical areas, especially with those areas outside of M1. While the basic connectivity of RFA has been described in a qualitative way, more quantitative details are needed to resolve this question. Due to the small size of the rat brain relative to larger non-human primate brains, injection cores have been relatively large in most tract-tracing studies, especially those conducted before the advent of labeled dextrans. Thus, maintaining tracer injections to a small region such as the RFA, occupying about 1 mm^2^ of the cortical surface (Barbay S et al. 2013), is challenging. As tract-tracing methods have evolved, smaller and more confined injection cores have allowed for a more precise picture of corticocortical connections. Few studies using more modern tracing techniques have been conducted in RFA, and none have used quantitative approaches. Lastly, although the laminar distribution of connectivity to/from RFA has been described in detail by examining labeling in coronal sections, tangential sectioning allows more precise definition of topographic relationships of terminal fields. The current study employed ICMS to define the boundaries of RFA, utilized small injections of labeled dextrans, and tangential sectioning to understand the anterograde and retrograde connectivity of this functionally important region of rat cortex.

The results of the current study provide a quantitative description of the relative density of connections of the RFA with other cortical areas. We confirm that the ipsilateral corticocortical connectivity of RFA is not diffuse throughout the cortex, but has clusters of connectivity with a preference for motor regions and higher order somatosensory processing areas. Further, we rank ordered the density of anterograde connections, providing further documentation of the corticocortical hierarchy. Retrograde labeling demonstrated a similar pattern, and suggests that corticocortical connectivity with RFA is largely reciprocal.

## Materials and Methods

Adult male Long-Evans hooded rats (n=8 entered study; n=6 analyzed; 370-450 g; 3-5 months of age Harlan, Indianapolis, IN) were singly-housed with a 12 hr:12 hr light:dark cycle. Food and water were provided *ad libitum*. The Institutional Animal Care and Use Committee of the University of Kansas Medical Center approved all animal use. All experiments were conducted in accordance with the guidelines published in the NIH *Guide for the Care and Use of Laboratory Animals*.

### Surgical Procedure

Isoflurane sedation was followed by ketamine [100-80 mg/kg, intramuscularly (IP)] and xylazine [30 mg/kg, intraperitoneal (IM)] anesthesia. Supplemental doses of ketamine (20 mg/kg IM) were provided throughout the procedure as needed to maintain stable anesthetic depth. After the rat was secured in a stereotaxic frame, Bupivacaine (2.5 mg, local anesthetic) was applied to the scalp. A homeothermic blanket system maintained physiological body temperature. The scalp was incised and reflected, and muscles attached to the temporal and occipital ridges were released. The cisterna magna was opened to relieve cerebrospinal fluid, and a craniotomy performed from +5 anterior to and -4 mm posterior to Bregma, and from +1 mm lateral to the midline to the temporal ridge. The dura was reflected and warm sterile silicone oil applied to the cortex.

Motor areas were identified by intracortical microstimulation (ICMS) methods (Urban ET, 3rd *et al*. 2012; Barbay S *et al*. 2013). Briefly, a digital photomicrograph of the cortical surface vasculature was obtained through the surgical microscope and overlaid with a grid pattern (250 µm) in image software (Canvas, Deneba Software, Miami, FL). A tapered and beveled glass micropipette (20 µm outside diameter) filled with concentrated saline solution (3.5 M), served as the microelectrode. The pipette was inserted 1725 µm below the cortical surface at every other grid intersection to provide a resolution of 500 µm. A stimulation pulse train (40 msec duration), consisting of 13 monophasic cathodal pulses (200 µsec duration, 350 Hz) was delivered each second from an electrically isolated, charge-balanced (capacitively-coupled), constant-current stimulation circuit (BSI-2, Bak Electronics Inc, Mount Airy, MD). The current was increased gradually from 0 µA until a movement was visible, then reduced until the movement was no longer visible. Stimulation of sites defined as “nonresponsive” did not elicit movements at the maximum current level of 60µA.

After defining the borders of RFA, a micropipette containing the neuronal tract tracer, biotinylated dextran amine, 10,000 MW (BDA10kDa, 10% w/v in 0.9% sterile saline) was placed at approximately the center of RFA. The injection needle included a tapered glass micropipette cut to 60µm outside diameter and was attached with beeswax to a 1µL Hamilton syringe (30100, Hamilton Company, Reno, NV). It was actuated by a microinjector (Micro4, World Precision Instruments, Sarasota, FL). Injection depth was controlled by a hydraulic Microdrive (650 Micropositioner, David Kopf Instruments, Tujunga, CA) on a stereotaxic arm. BDA10kDa was pressure injected in 3 boluses of 33.3nL each (100nL total) at 1500, 1250 and 1000µm below the cortical surface; CTB- 647 injections were delivered in 2 boluses of 75nL each (150nL total) at 1500 and 1000µm below the cortical surface. Cholera toxin beta subunit conjugated to AlexaFluor 647 (CTB647, 5 µg/µL in 0.9% sterile saline, C34778, Invitrogen, Grand Island, NY) was injected (with the same configuration and outside diameter as BDA10kDa) at 2 sites that were each roughly 1mm caudal to the caudal border of CFA as defined by ICMS, serving as fiducial markers.

The cortical surface was rinsed with warm sterile saline (0.9%) and covered with a silicone sheet (Invotec International Inc, Jacksonville, FL), gel foam, (Surgifoam, Ethicon, Sommerville NJ) and dental acrylic and resin (Lang Dental Mfg Co Inc, Wheeling, IL) to form a protective cap over the craniotomy. The skin was sutured, penicillin injected (45,000 U, SQ) into the nape of the neck and local anesthetic (Bupivicaine, 2.5mg, APP Pharmaceuticals, Schaumburg, IL) and topical antibiotic (Vetropolylycine gel, Dechra Veterinary Products, OP, KS) applied. Buprenorphine (0.05 mg/kg SQ, Reckitt Benckiser Farmaceuticals Inc, Richmond, VA) and acetaminophen (40 mg/kg oral) were given after the surgery for pain management. The rat was allowed to recover on the heating pad until it was alert and moving spontaneously, and then returned to its home cage. Three additional doses of buprenorphine and acetaminophen were given during the subsequent 48 hours.

### Histology

#### Tissue harvesting

Seven days after the surgical procedure, rats were sedated with isoflurane, and euthanized with Beuthenasia-D (390mg pentobarbital, 50mg phenytoin sodium IP, Shering Plough Animal Health, Union, NJ). After thoracotomy and rib cage reflection, heparin sodium (500 USP Units, Hospira Inc, IL) was injected into the left ventricle, and exsanguination was achieved through transcardial perfusion of saline solution [0.9% saline in distilled water, heparin sodium (1,000 USP Units, APP Pharmaceuticals, Schaumburg, IL) and lidocaine HCl (20 mg, APP Pharmaceuticals, Lake Forest, IL)] followed by 3% paraformaldehyde in 0.9% saline. The brain was extracted, both hemispheres of cortex were separated from the underlying structures, and flattened between glass slides. The flattened cortices were exposed to 4% paraformaldehyde-20% glycerol in 0.9% saline (2 hr), 20% glycerol-2% dimethylsulfoxide in 0.9% saline (overnight), and 20% glycerol in 0.9% saline (24 hr). Each flattened cortex was sectioned at 50 µm thickness on a freezing microtome chilled with dry ice. Individual sections were placed in 0.1 M PBS solution and maintained at 4°C.

#### Cytochrome oxidase staining

Cytochrome oxidase staining was used in selected sections to aid in the delineation of cortical sensory areas. Sections were placed in 0.1 M PBS solution and allowed to float. Then, sections were inspected with the unaided eye for the S1 representation, which is visible as several slightly opaque white areas within the translucent section. Sections with the most complete representations (3 to 4 sections/cortex) were chosen for cytochrome oxidase (CO) staining. After rinsing (2 x 10 min in 0.1 M PBS), floating sections were reacted with CO solution at 37°C containing cytochrome c oxidase (20 mg, Sigma, #C2506-500MG), sucrose (4 g, Fisher Scientific), and DAB (50 mg) per 100 mL of 0.1 M phosphate buffered distilled water (pH 7.4). Sections were allowed to react for ∼2-3 hr until dark CO-rich areas were easily detectable against the lighter background. The sections were then rinsed (2 x 10 min) in 0.1 M PBS.

#### BDA10kDa visualization

All sections underwent a standard staining procedure using Avidin-Biotin Complex (ABC) linked to peroxidase with 3,3’ Diaminobenzidine (DAB, MP Biomedicals, Solon, OH, #980681) reaction product as the chromogen. Sections were rinsed in 0.1 M PBS (2 x 10 min with agitation), then exposed to 0.4% Triton X-100 (Sigma, #X100-500ML) in 0.05 M PBS (1 hr with agitation). Sections were rinsed in 0.1 M PBS (3 x 10 min, with agitation). Sections were incubated overnight in 0.1 M PBS with reagents “A” and “B” added according to Vectastain Elite Kit (Vector Laboratories, Burlingame, CA, #PK6100). Then, sections were rinsed (4 x 10 min, 0.1 M PBS), and exposed to DAB solution (0.05% w/v DAB and 0.01% v/v H2O2 in 0.1 M PBS). Sections were wet mounted in 0.05 M PBS onto subbed slides and allowed to dry overnight.

#### BDA10kDa signal intensification

Sections on slides were dehydrated in ascending alcohol concentrations (50%, 70%, 95% and 100% for 5 min each), cleared with xylene (5 min), then rehydrated by reverse order of alcohol concentrations. Sections were exposed to 1.42% silver nitrate in distilled water (55°C, 1 hr), rinsed (15 min, distilled water), exposed to 0.2% gold chloride (10 min), rinsed (15 min, distilled water), exposed to sodium thiosulfate (5 min), and rinsed (15 min, distilled water). Finally, sections were dehydrated again, as described above, cleared in xylene, and coverslipped with DPX mounting medium (Sigma, #44581-500ML).

### Stereological Analysis

#### Terminal bouton and soma quantification

Section outlines were traced with the aid of a computerized microscope with motorized stage (Axiophot 2, Zeiss) and stereology program (Stereoinvestigator, Microbrightfield). Because of the large number of regions of interest, we used a method previously developed for a survey of premotor cortex connections in squirrel monkey (Dancause N, S Barbay, SB Frost, EJ Plautz, et al. 2006). A 100 µm square grid was overlaid on the flattened cortical outline. As the section thickness was 50 µm, voxels with dimensions 100 x 100 x 50 µm were examined systematically throughout the section. Acronyms used for the regions of interest are defined in Table 1.

**Table 1:** Abbreviations

|  |  |
| --- | --- |
| AGm | Medial agranular cortex |
| ALBSF | Anterolateral Barrel Subfield of S1 |
| Aud | Auditory Cortex |
| BDA | Biotinylated Dextran Amine |
| CFA | Caudal Forelimb Area |
| CFAc | Caudal Forelimb Area, caudal portion |
| CFAov | Caudal Forelimb Area, overlap zone with S1 |
| CFAr | Caudal Forelimb Area, rostral portion |
| CO | Cytochrome Oxidase |
| CTB | Cholera Toxin Beta subunit |
| DAB | Diaminobenzidine |
| FL-ICMS | Forelimb ICMS Area |
| FR | Frontal Cortical Area |
| FRo | Frontal Cortical Area, other |
| GZ | Granular Zone of S1 |
| GZo | Granular Zone of S1, other |
| ICMS | Intracortical Microstimulation |
| IN | Insular Area |
| DZ | Dysgranular Zone of S1 |
| DZo | Dysgranular Zone of S1, other |
| M1 | Primary Motor Cortex |
| PirOI | Piriform and Olfactory Cortex |
| PMBSF | Posteromedial Barrel Subfield of S1 |
| PPCI | Posterior parietal cortex, lateral aspect |
| PPCm | Posterior parietal cortex, medial aspect |
| PR | Parietal rhinal cortex |
| PRh | Perirhinal Cortex |
| PV | Parietal Ventral Area |
| RFA | Rostral Forelimb Area |
| RS | Retrosplenial Area |
| S1 | Primary Somatosensory Area |
| S1<sub>FP</sub> | Forepaw Area of S1 |
| S1<sub>FPO</sub> | Forepaw Area of S1, other |
| S2 | Second Somatosensory Area |
| TP | Temporal Posterior Cortex |
| Vis | Visual Cortex |

Axons labeled with BDA10kDa appeared as dark lines with varicosities or boutons (Fig. 1E). A bouton was defined as a dark (chromogen dense), round object about twice as wide as the thin dark fiber on either side of it (*en passant* bouton), or at the end of a thin projection off the main axon (terminal bouton). Boutons were counted and recorded semi-quantitatively within each 100 µm x 100 µm x 50 µm (section thickness) voxel. Voxels containing 2-30 boutons were marked with a blue dot (Fig. 1I), and voxels containing greater than 30 boutons were marked with a red dot (Fig. 1l).

**Figure 1.**
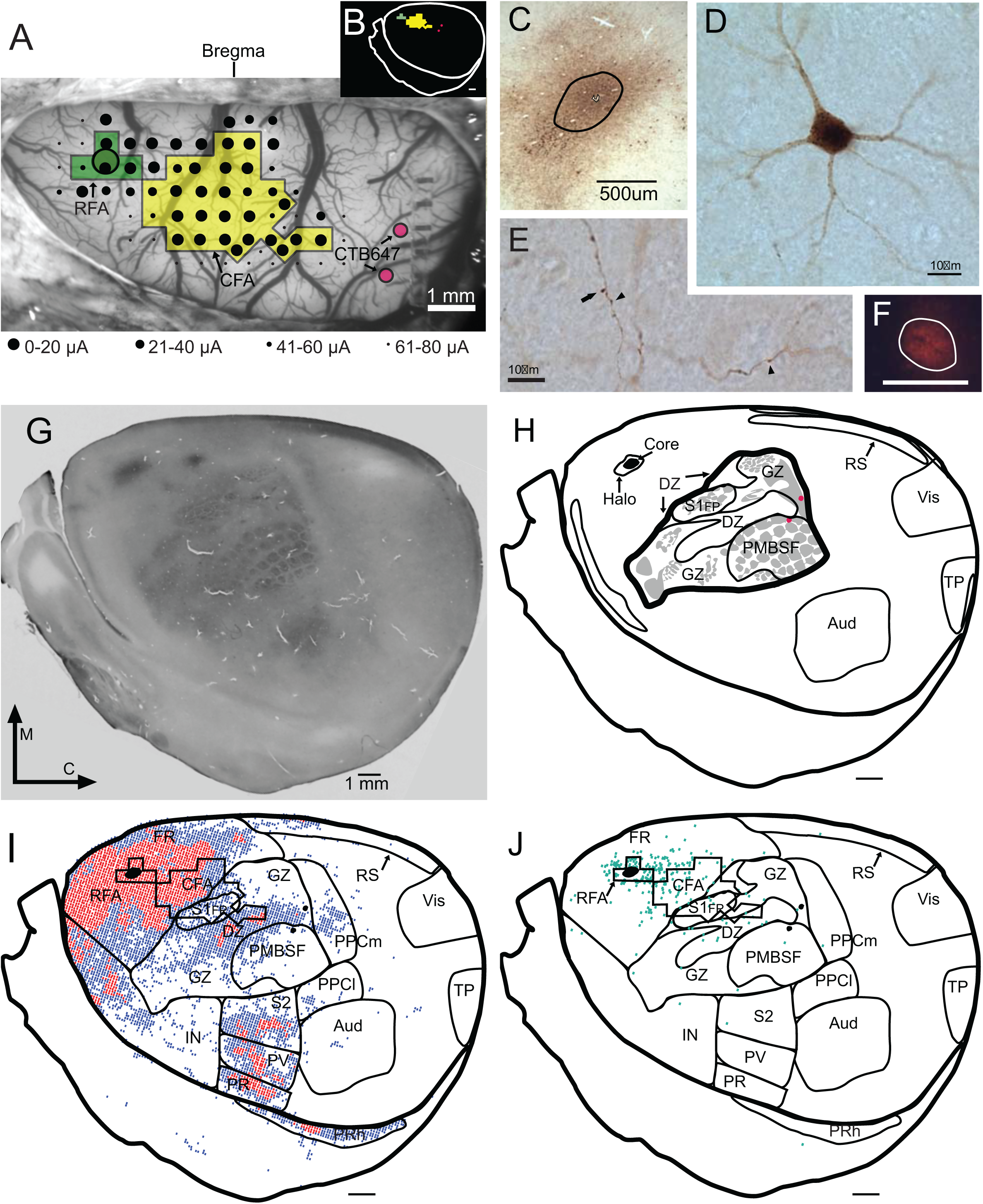
Experimental overview. **A**. Photomicrograph of the cortex surface vasculature following craniotomy (case R11-34). Location of ICMS-defined RFA (green) and CFA (yellow) superimposed on surface vasculature. Black dots indicate microelectrode penetration sites. The size of the dots is inversely related to current threshold for evoking movements (see legend). Circle within RFA indicates site of BDA injection. Red dots indicate site of CTB647 injections used as fiducial markers. **B**. Inset to show relationship of RFA/CFA locations to sections in G-J. **C**. Photomicrograph of BDA injection core (black outline) within RFA in 50 µm thick, flattened, tangential histological section. **D**. Retrogradely- labeled neuronal soma and processes stained with BDA (brown). **E**. BDA-labeled axonal projections with boutons (arrow heads). **F.** Photomicrograph of CTB647 injection site. Scale bar = 0.1 mm. **G**. Flattened, tangential section stained with cytochrome oxidase to indicate location of sensory areas. Somatosensory representations are outlined. **H**. Outline of sensory and other relevant areas based on cytochrome oxidase and BDA staining in G. The granular zone of S1 (light grey areas) is further divided into S1H, GZ, and PMBSF. **I**. Location of 100 µm X 100 µm (plotting area) x 50 µm (section thickness) voxels containing BDA- labeled boutons. Blue dot = voxel containing 2-30 boutons; red dot = voxel containing >30 boutons. **J**. Location of voxels containing BDA-labeled somata (green dots). Abbreviations: RFA = rostral forelimb area; CFA = caudal forelimb area; M = medial; C = caudal; Aud = primary auditory cortex; GZ = granular zone of S1; DZ = dysgranular zone of S1; PMBSF = posteriomedial barrel subfield of S1; RS = retrosplenial area; S1FP = forepaw area of S1; TP = temporoposterior cortex; V1 = primary visual cortex.

Although BDA10kDa is an effective anterograde tract tracer, retrograde labeling occurs as well. BDA-labeled neuronal soma were plotted and counted on the same section as boutons. A neuronal soma was defined as a uniformly dark, smooth-edged shape with evidence of at least one thin projection emanating from it (Dancause N, S Barbay, SB Frost, EV Zoubina, et al. 2006) (Fig 1D). BDA-labeled soma were marked with a green dot (Fig 1J).

Within a given region, the density of boutons was greatest in deeper cortical laminae and least in superficial laminae. However, the distribution of BDA-labeled boutons and soma was similar throughout cortical depths, similar to descriptions in previous connectivity studies in rat (Reep RL et al. 1987; Rouiller EM *et al*. 1993). The current experiment used a superficial layer section in order to more easily delineate between regions of interest, as deep layers have a larger more diffuse projection pattern. Thus, although variations in overall density exist from superficial to deep laminae, it was deemed that a single section was representative of a particular animal’s connectivity patterns. Due to the extensive amount of time required to plot BDA-labeling in each section, this was the most feasible approach to describing consistent connectivity patterns in the current sample.

To verify the validity of this approach for estimating voxel densities, we compared values with unbiased stereological techniques within eight regions of interest throughout all available sections in a random sample of three of the six rats. Estimations of terminal boutons in the eight regions were performed using the stereological probe, the Optical Fractionator (Microbrightfield, Colchester, VT), according to methods described previously (Gundersen HJ and EB Jensen 1987; Gundersen HJ et al. 1988). Briefly, the Optical Fractionator is a stereological probe that estimates population sizes by counting objects with optical disectors in a random, systematic sample within a volume of tissue located within a region of interest (ROI) (West MJ et al. 1991). This allowed us to test the validity of the voxel counts without performing unbiased stereology on every area in every animal. Spearman’s rho was used to compare ranks between the two approaches.

#### Alignment procedure

Fiducial markers were used to align the ICMS maps from the surgical procedure with the section outlines, voxel counts, neuronal soma counts, and CO-rich zones drawn in StereoInvestigator. The location of two injections of CTB647 and 1 injection of BDA10kDa injection cores were marked on all ICMS maps and section outlines. Section outlines and ICMS maps were overlaid in Photoshop, and the ICMS map was scaled and rotated until the 3 injection cores were in register. Numerous symbols used to designate voxels (red and blue dots) were present in the region of RFA and FL-ICMS. Careful attention was paid to the relationship between the borders of RFA and FL-ICMS to the symbols, which helped maintain correct proportion while transferring the RFA and FL-ICMS outlines into StereoInvestigator.

After visual inspection of the sectioned tissue, sections 8 through 24 were determined to be consistently intact in all animals and were used for further analysis. Every other section (nine sections per animal) was used for determining BDA injection core size. Both BDA and CTB647 (Fig. 1F) injection cores were outlined on sections. The section outlines and the ICMS maps were aligned using these cores as fiducial markers.

#### Calculation of cortical surface area for regions of interest

The areal representations of the regions of interest (as determined by alignment of CO stained sections, ICMS maps and voxel clusters) were outlined in StereoInvestigator, and imported into a graphics software program (Illustrator, Adobe). The area (mm^2^) of each region was then measured with Image J (NIH).

### Statistical Analysis

Both voxel counts and soma counts were normalized to the surface area of the region of interest. These values (labeled voxels/mm^2^, and somata/mm^2^) were examined using a statistical program (JMP v10, SAS Institute). Since the variance in the different regions was determined to be unequal (O’Brien test, F = 4.34; p < 0.0001), a nonparametric analysis, Wilcoxon Rank Sum test, was used to compare values in the different regions. Z-scores generated by the Wilcoxon test were then used to define cutoff levels for dense, moderate, sparse and negligible connectivity.

### Region Nomenclature and Identification

The nomenclature, location and description of the regions of interest utilized in this study are derived from various sources to achieve the most accurate and reliable description. Regions were identified using a variety of criteria including, response to ICMS, CO staining, obvious clustering of voxels, or spatial relationship to other regions, and are consistent with previously reported nomenclature.

## Results

Of the eight animals that underwent surgical procedures, six were used for examination of the distribution and quantification of BDA-labeling in neuronal somata and terminal boutons. One animal died at the end of the surgical procedure. In another animal, the CO staining procedure failed to reveal the S1 representation.

In each of the six animals used for analysis, BDA-labeling of terminal boutons and neuronal somata was clearly visible in various regions of the cortex. In the sections that follow, first the regions of interest are defined. Next, the spread of the BDA injection cores are compared to the neurophysiologically- defined RFA borders. Then, the distribution of BDA-labeled terminal boutons and neuronal somata is described with respect to the regions of interest. Finally, the quantitative results (relative density of BDA-labeled terminal boutons and neuronal somata) are described.

### Identification of Regions-of-Interest

A total of 22 regions of interest were identified based on a combination of features, including ICMS mapping data, cytochrome oxidase labeling, clustering of BDA labeling, relative topographical location, and neurophysiological/neuroanatomical overlap. The tangential sectioning of the cortex aided in the alignment of the multiple data sets, and ultimately in the standardization of nomenclature across the sample of animals. Since some of the 22 regions of interest overlapped broader designations (e.g., FRo, GZo, DZo and S1FPo) are used here to define mutually exclusive territories for quantitative analysis. For example, GZo refers to the cortical area corresponding to the granular zone of S1 (GZ), but excludes GZ cortex overlapping with CFA. GZo can be read “Granular Zone, other”.

#### Identification of CFA and RFA based on ICMS-evoked movements

ICMS maps of evoked movements were successfully obtained in each of the six animals used for analysis. Movements of the forelimb were evoked by ICMS within two clusters corresponding to the typical locations of the RFA and CFA using currents ≤ 60µA. (Neafsey EJ and C Sievert 1982; Nishibe M et al. 2010). Clusters of sites at which ICMS evoked forelimb movements were surrounded by sites at which ICMS evoked neck, jaw, orofacial and vibrissae movements, or sites at which ICMS evoked no movement. The smaller cluster corresponding to RFA was located between +3.7 and +2.7 mm anterior to Bregma and 2.0 to 4.0 mm lateral to the sagittal suture (Fig. 1A). The larger cluster corresponding to CFA primarily was located between +2.7 anterior and -1 mm posterior to Bregma and 2.0 to 4.5 mm lateral of the sagittal suture. In some cases, a narrow, caudal extension of the CFA movement map was observed. Since CFA was defined by the area from which forelimb movements could be evoked by ICMS currents ≤60 µA, we included this caudal extension as part of CFA.

#### Regions of interest identified by cytochrome oxidase staining

Cytochrome oxidase staining revealed darkly-stained zones within the granular cortex of flattened sections (Fig. 1G). These CO-rich zones were useful in identifying the primary sensory areas, S1, Vis, and Aud, as well as RS and TP cortex. CO staining was especially useful for identifying the prominent barrel field (PMBSF) within S1. S1 is a critical landmark in aligning data sets. These areas are consistent with Remple et al. 2003 (Remple MS et al. 2003) Chapin and Lin 1984 (Chapin JK and C-S Lin 1984).

Vis and TP are CO-dense regions coursing from the caudal edge of the cortex in a wide triangular shape. Both regions are thinnest at the rostral vertex. Vis is consistent with the pooled Oc1M and Oc1B of Zilles (Zilles KJ 1985). TP is the smaller triangular region directly lateral to and separated from Vis by a CO-sparse strip of cortex, consistent with Krubitzer (Krubitzer L et al. 2011). Aud is a large circular CO-dense region lateral and caudal to the S1, consistent with Au of Remple et al. (Remple MS *et al*. 2003), and the 5 auditory fields of Polley (Polley DB et al. 2007).

RS comprises a thin CO-dense region coursing along the medial edge of the caudal half of the flattened cortex (Harley CA and CH Bielajew 1992). The rostral aspect of RS stops at the caudal border of FR; the lateral border is concurrent with the CAA; the caudal aspect ends 1-2 mm from Vis; the medial border extends to the medial border of the cortex.

#### Identification of regions of interest based on clusters of BDA-labeling

The locations of discrete clusters of voxels containing BDA-labeled terminal boutons superimposed on section outlines led to the identification of several regions of interest. S2, PV, PR, PPCm and PPCl, as defined here, are in accordance with Remple et al. (Remple MS *et al*. 2003), and Fabri and Burton (Fabri M and H Burton 1991), though PPCm and PPCl are sometimes called PM and PL, respectively. Discrete dense clusters of labeling were identifiable lateral to S1, which spanned the area directly lateral to the caudal half of ALBSF and the rostral half of PMBSF. Proceeding from medial to lateral, S2, PV and PR occupy the cortex lateral to S1, i.e., above the rhinal fissure, rostral to Aud and caudal to IN. These three areas each have generally the same dimensions, 1-3 mm in the mediolateral and 2-3 mm in the rostrocaudal dimensions. As the boundaries of labeled vs. unlabeled tissue were relatively sharp, the extent of the regions was determined by the largest extent of the cluster of voxels with labeled boutons.

PPCm and PPCl were identified by small clusters of voxels with labeled boutons, which were located caudal to and caudolateral to S1, respectively. PPCm shares its rostral border with S1. The extent of the labeled area was ∼1.2 mm anterior to posterior. PPCl shares a border with PPCm and S1 medially, S2 rostrally, and Aud laterally. This distribution pattern is similar to those reported in other published reports (Fabri M and H Burton 1991; Reep RL and JV Corwin 2009).

#### Identification of regions of interest by topographic relationships to other identified regions

After aligning CO-rich regions with the distribution of BDA-labeling, additional regions were identified by their spatial relationship to other regions. FRo and IN were largely identified by their relationship to S1. FR occupies the rostral half of the medial cortex as well as the frontal pole, bordering S1 along its lateral caudal aspect. The definitions of FR and IN cortex follow Zilles 1985 (Zilles KJ 1985) with some modification. All of the Frontal regions of Zilles (Fr1, Fr2, and Fr3, and Cingulate cortex) were pooled into one region: FR. The term FRo is used here to distinguish the frontal rostral area that excludes RFA, the focus of the study. Zilles’ insular cortex (AID, AIV) was pooled with the rostral half of Vi to comprise IN in the present report. IN occupies the lateral aspect of the cortex and is bordered by FR and S1 at its medial aspect, shares its caudal border with S2, PV and PR, and the rhinal fissure laterally. The border between IN and FR was drawn from the rostromedial edge of S1 to a small consistent area of dense axons and boutons in the rostral pole.

#### Identification of regions of interest based on neurophysiological/ neuroanatomical overlap

Some regions were identified through the overlap of ICMS mapping data with CO-rich areas. These overlapping zones were treated as separate regions in the quantitative analysis. Most importantly, several investigators have noted the overlap zone between CFA and S1FP, i.e., the forelimb portion of GZ(Sanderson KJ et al. 1984; Tennant KA et al. 2011). Due to its potential functional importance in interpreting RFA connectivity, the CFA was divided into three partitions (Fig. 2). CFAr was designated as the cortex that includes the rostral portion of CFA, i.e., the portion of CFA anterior to the overlap zone with S1FP. Thus, CFAr overlaps with the rostral portion of DZ and the caudal portion of FR. CFAov is the CFA overlap zone with GZ, including S1FP. CFAc was designated as the cortex that includes the caudalmost portion of CFA, overlapping with a portion of DZ.

**Figure 2.**
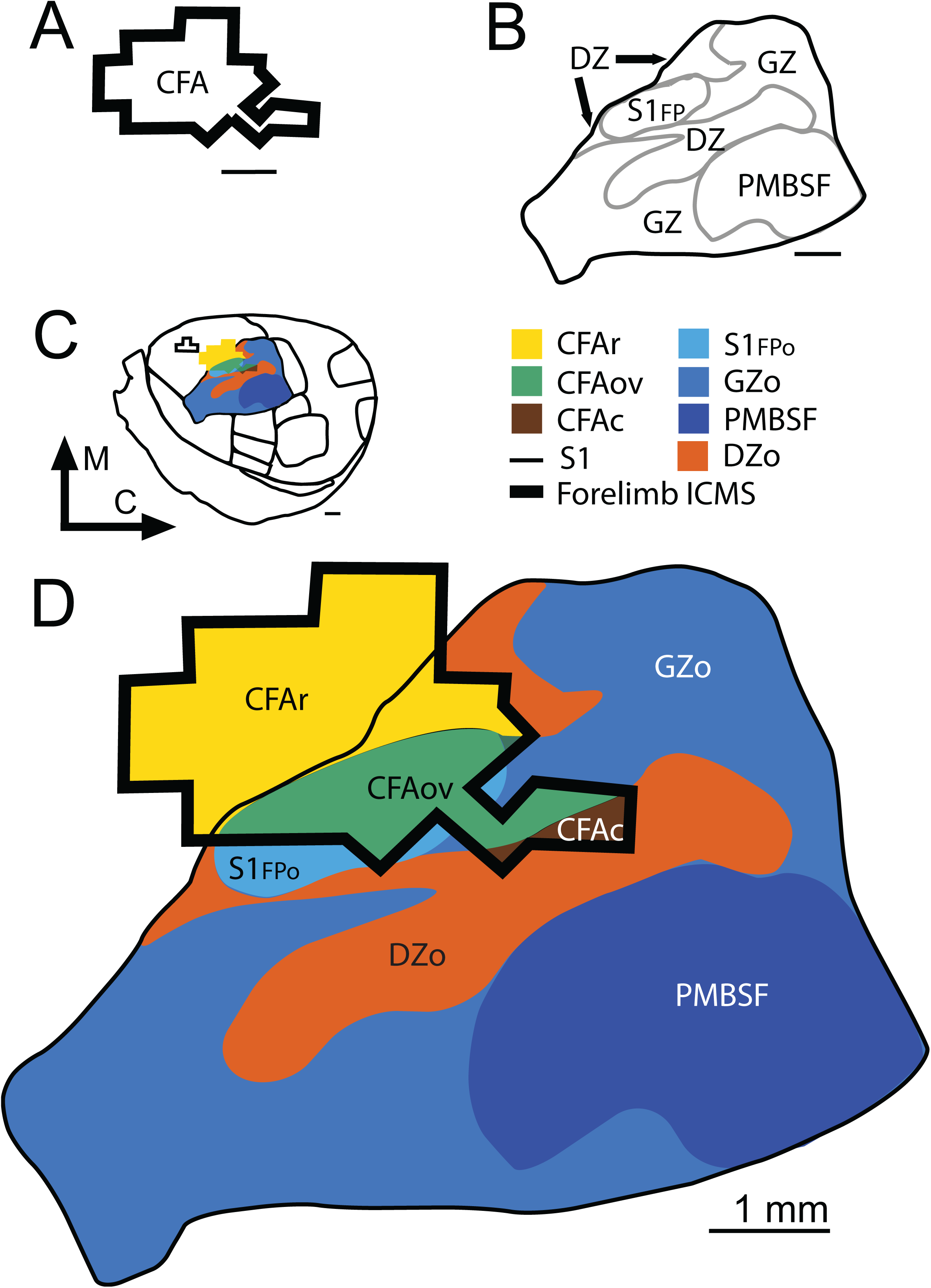
Overlap of CFA and S1. **A**. Caudal forelimb (motor) area as defined by ICMS. **B**. S1 subdivisions as defined by neurophysiological recordings and cytochrome oxidase staining. **C**. Location of cortical areas on flattened, tangential section. **D.** Enlarged view of ICMS-defined forelimb motor area and S1 subdivisions. Abbreviations: FL-ICMS = ICMS-defined caudal forelimb motor area; CFAr = rostral portion of CFA; CFAov = overlap zone between CFA and S1FP; CFAc = caudal portion of CFA; S1FP = forepaw somatosensory area. Note that the entire forepaw somatosensory area in D comprises S1FP and CFAov.

With respect to the histologically-defined S1, the size and location of motor representations in CFAr and CFAov was similar across animals. CFAr was located along ∼2 mm of the rostrolateral border of S1FP, extending rostrally ∼2 mm from the S1FP border. In all six animals, CFAov was co-extensive with S1FP, arranged in an oblique angle along the rostral aspect of S1, and extended more caudally into non-FP areas of GZ. Its dimensions were ∼2 x 3 mm. In contrast, there was variability in the extreme caudal extent of CFA. In four of the six animals, ICMS at low current levels (< 60 µA) evoked forelimb movements in regions caudal to GZ (Fig. 3B, C, E, and F). This region extended caudally as a narrow strip of one to four sites at which ICMS evoked forelimb movements. The caudal-most sites overlapped with the histologically defined DZ (Chapin JK and C-S Lin 1984). CFAc did not overlap PMBSF in any of the six animals. In the four animals in which CFAc was identified, forelimb movements were evoked at a total of 15 sites. Of these 15 sites, wrist extension was evoked at 13 sites, while finger flexion was evoked at one site, and elbow extension at one site.

**Figure 3.**
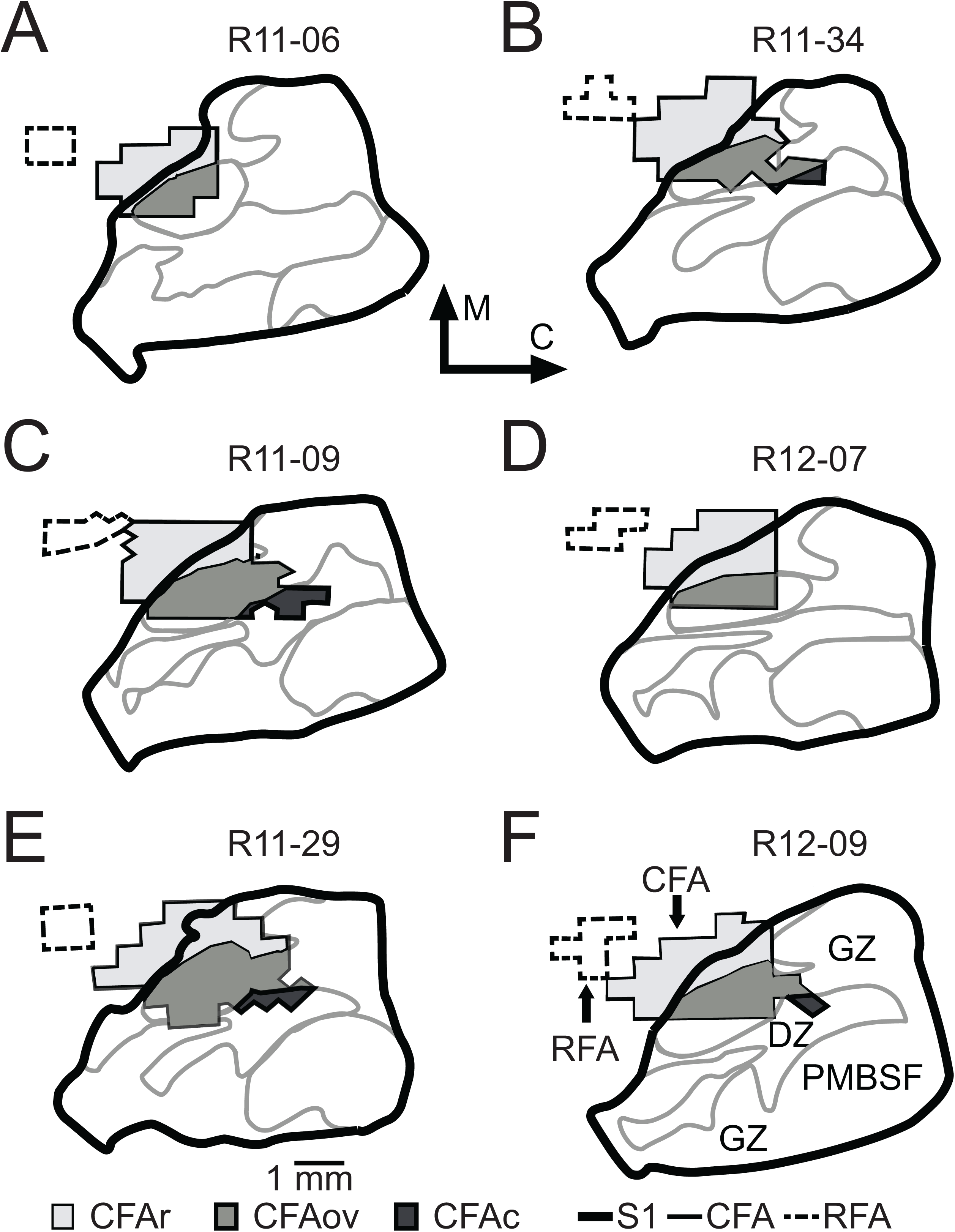
Overlap of CFA and S1 in individual cases.

RFA formed a segregated cluster of sites where ICMS evoked movements of the forelimb, as described in a previous section. RFA did not overlap with any of the CO-positive regions.

### Extent of BDA Injections

In the flattened, tangentially-sectioned cortex, the BDA injection core was defined as the area of dark, uniformly-speckled staining with little identifiable cellular structure. The halo was identified as a larger area of dark staining extending beyond the core. While indistinguishable at low magnification, at high magnification, a large number of BDA-labeled axons could be observed immediately around the injection core (Fig. 1C). This region was defined as the halo. The injection core identified in nine BDA-labeled sections in each animal was aligned using fiducial landmarks and superimposed in the neurophysiological data. In each of the six cases, the BDA injection core was within the borders of RFA as defined by ICMS (Fig. 4).

**Figure 4.**
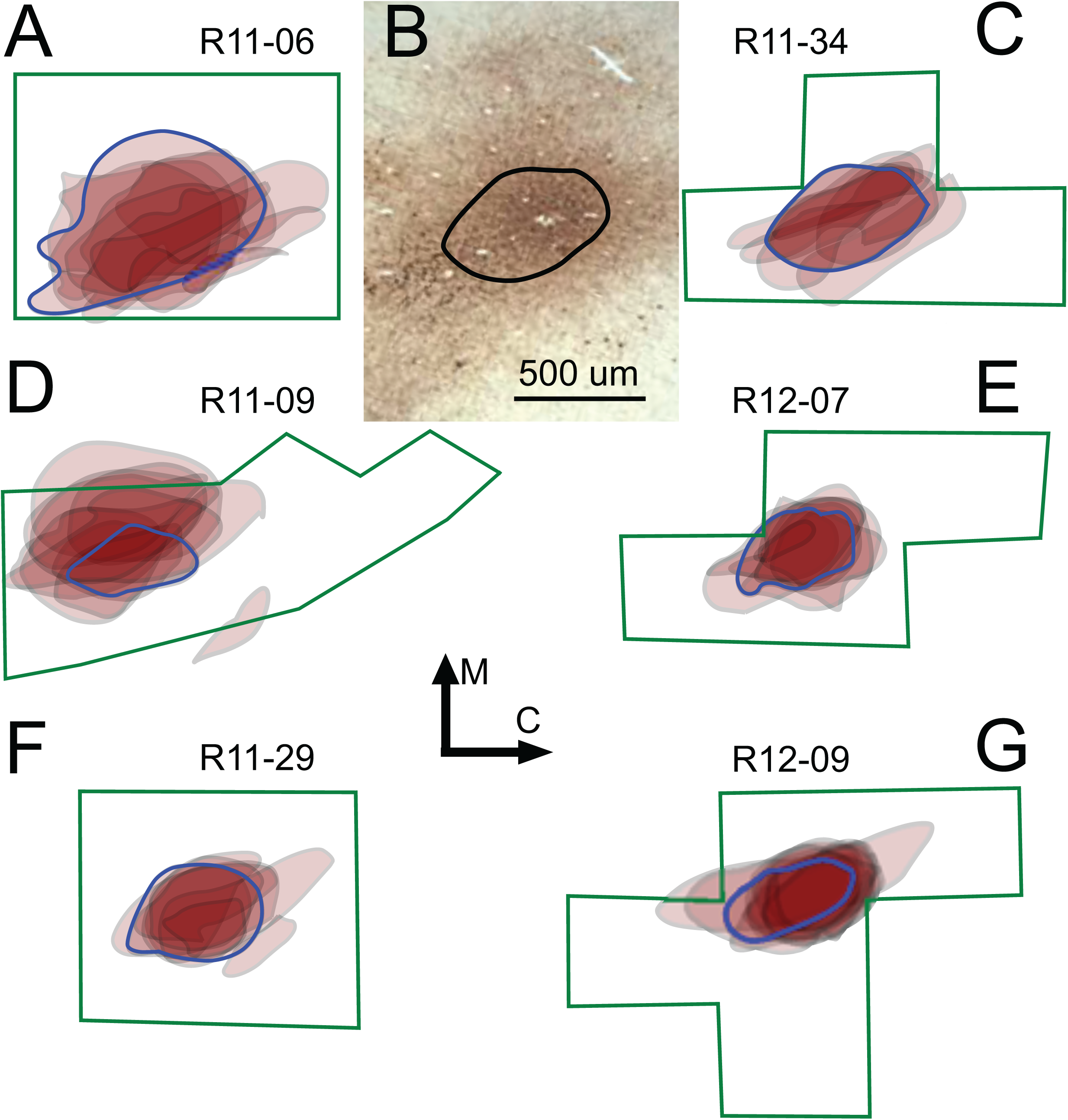
Extent of BDA injection cores relative to ICMS-defined RFA extent. In **A** and **C-G**, individual cases are illustrated. For each case, each tangential section was aligned and the vertically-aligned BDA injection cores. Thus, darker areas represent a greater degree of overlap with other sections in the case. The injection core outline from the section used for bouton (voxel) counting is outlined in blue. Note that the BDA injection cores are largely confined to the ICMS-defined CFA territory. **B**. Photomicrograph of representative BDA injection core (black line). See methods for definition of core. All sections are scaled to the scale bar shown in **B**.

### Qualitative Distribution of BDA-labeled Boutons

Despite the relatively restricted injection cores, in each of the six animals, large numbers of BDA-labeled boutons were visible throughout frontal, parietal, insular and peri-rhinal cortex. Distributions were qualitatively similar in different animals, despite differences in total numbers of labeled boutons. As described in methods, each 100 x 100 x 50 µm voxel throughout a single section through the middle layer of cortex was examined, and coded separately if there were 0-1 (no color code), 2-30 (blue), or >30 (red) boutons per voxel. The distribution of voxels with labeled boutons (hereafter simply called “voxels”) in the animal with the fewest voxels (R11-29; Fig. 5A-B), and the animal with the most voxels (R11-09; Fig. 5C-D), is shown in Fig. 5 relative to the region-of-interest borders. Regardless of differences in total number, the relative distribution was similar between animals. Numerous red (> 30 boutons/voxel) and blue (2- 30boutons/voxel) voxels were consistently found in FRo, RFA, CFAr, CFAov, DZo, rostral IN, S2, PV, and PR. Voxels present within GZ were confined to the rostral border centered around CFAov. Smaller numbers of primarily blue voxels were found in GZo, PMBSF, PPCm, PPCl, PRh and caudal IN. Relatively few scattered blue voxels were found within Aud, PirOl, Vis, TP and RS.

**Figure 5.**
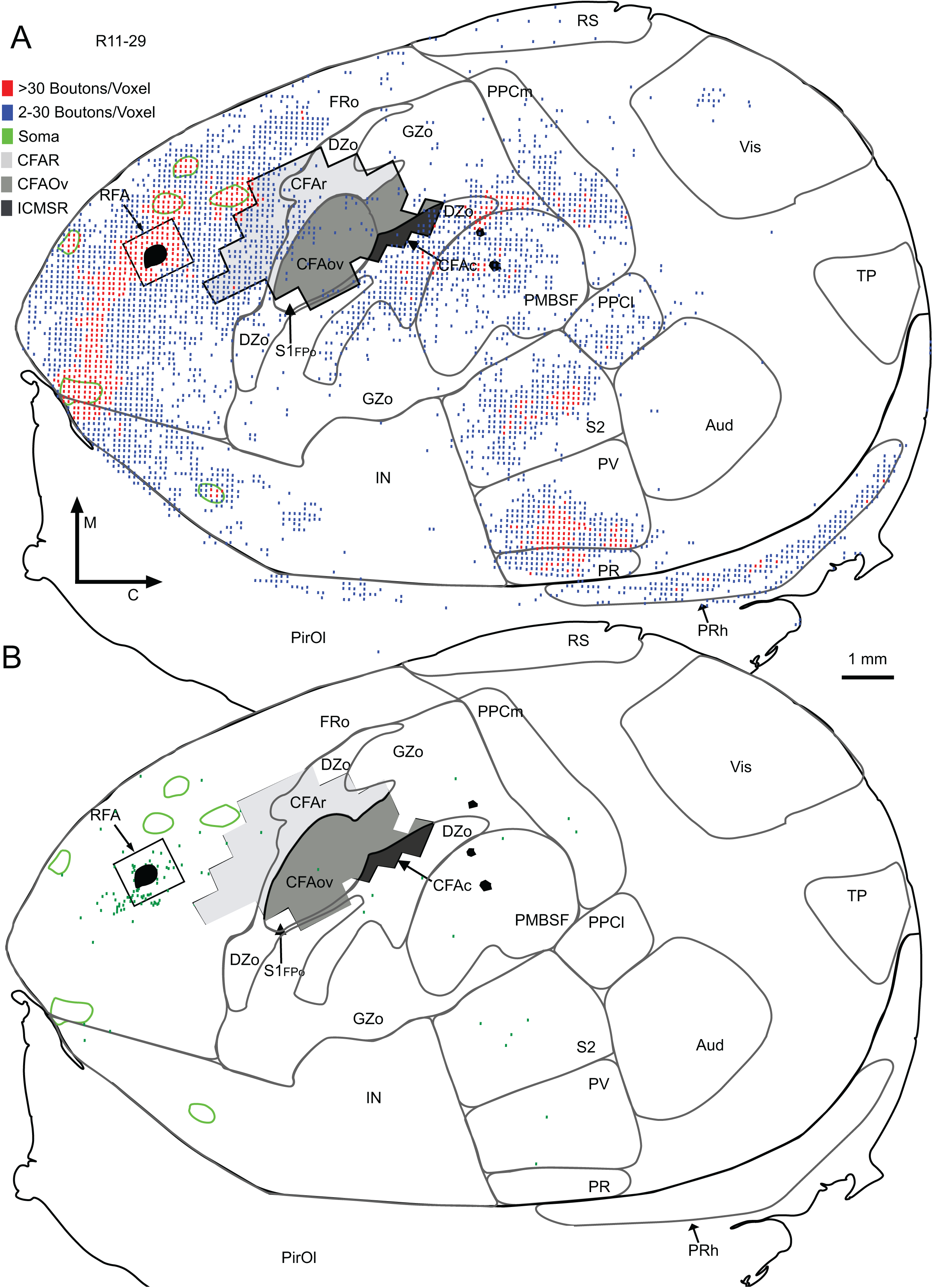
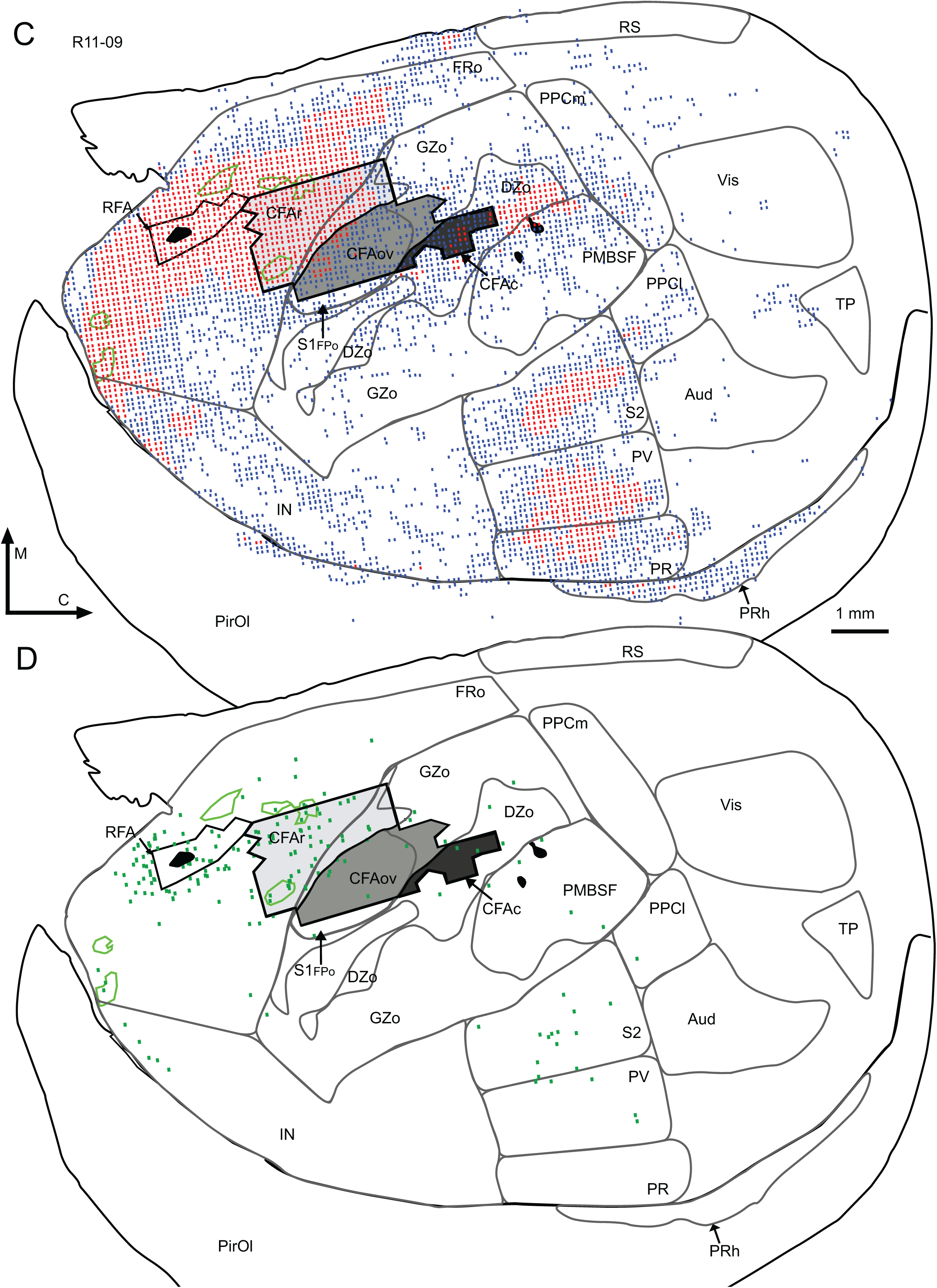
Spatial distribution of voxels with labeled boutons and labeled somata following BDA injection in RFA. **A** & **B** illustrate case R11-29, the case with the least number of labeled voxels; **C** & **D** illustrate case R11-09, the case with the most number of labeled voxels. Boutons (voxels) are shown in **A**, and somata in **B**. BDA injection core is shown as a filled black region within RFA. CTB647 injection site is shown as smaller black regions. Regions of unusually high bouton density are enclosed by green lines. A single, 50µm thick, tangential section in the middle layers of cortex is shown. In case R11-29 (**A**), large numbers of voxels with >30 BDA-labeled boutons (red dots) were observed in FR, S2, PV, PR, DZo, GZo, PM, rostral IN and PRh. Due to the 10kDa BDA used in these studies, relatively few BDA-labeled somata were observed in this case. In case R11-09 (**C**), large numbers of voxels with >30 BDA-labeled boutons (red dots) were observed in FRo, CFAr, the rostral part of CFAov, S2, PV, PR, DZ, extreme rostral PMBSF and rostral IN. More labeled somata were observed in this case (**D**), but the distribution was similar to that seen in (**B**). Qualitatively, the distribution was similar across cases.

Due to the perspective of the tangential sectioning, a clear distinction was found in S1. The caudal half of DZo was coextensive with an area of numerous voxels within the center of S1. The remainder of S1 (GZo, PMBSF and S1FPo) contained relatively few voxels. This pattern was evident in each of the six rats.

### Qualitative Description of BDA-labeled Somata

The distribution of retrogradely labeled somata paralleled the pattern of voxel distribution (Fig. 1 and 5). Sparsely distributed, labeled neuronal somata were distributed throughout the section with minimal clustering. Numerous labeled somata were found within RFA, FRo, CFAr, CFAOv, CFAr and DZ.

Relatively smaller numbers of labeled somata were found in PMBSF, GZo, S1FPo, S2, PV, PR, and PPCm. Labeled somata were rare in PPCl, RS, PirOl, and Vis, and were absent in PRh, Aud, and TP.

### Quantitative Description of Voxels with BDA-labeled Boutons

Total labeled voxels in each region, and voxels normalized to the region area, are shown in Table 2 and Figure 6, respectively. To verify the validity of this approach for rank-ordering densities, eight of the regions of interest (CFAr, PPCm, PV, CFAov, PPCl, PR, S2 and PMBSF) were selected to compare voxels/mm^2^ with estimates of bouton numbers using unbiased stereological techniques. Spearman’s rho nonparametric correlation analysis demonstrated a significant relationship (ρ = 0.6678; p = 0.0004), suggesting that the voxel approach using a single section for each case was valid for estimating relative densities of connections across animals.

**Figure 6.**
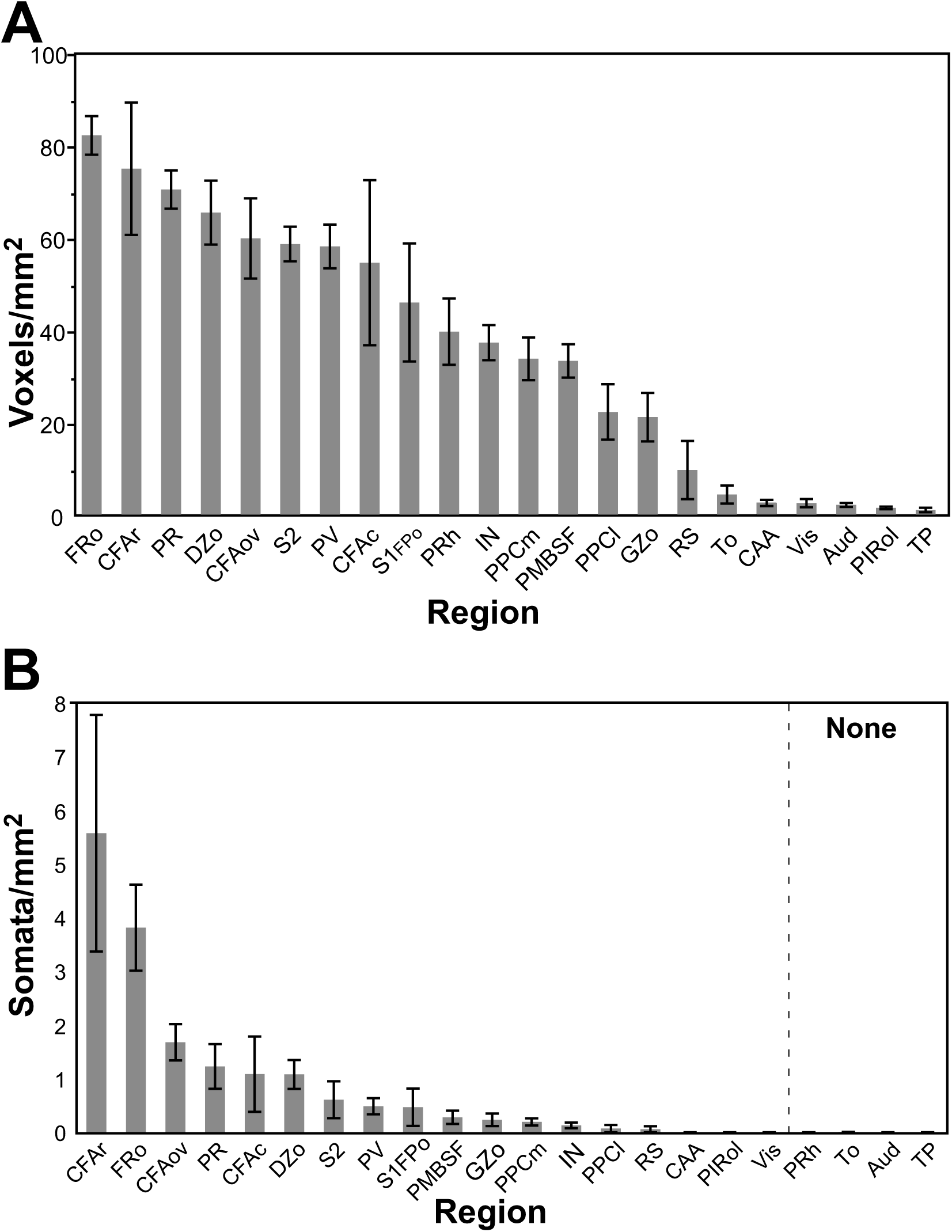
Quantitative distribution of BDA-labeled boutons and somata (normalized for region area) following BDA injection RFA. **A**. Mean number of voxels with ≥ 2 BDA-labeled boutons per mm^2^. **B**. Mean number of BDA-labeled somata per mm^2^.

**Table 2.** Means and Standard Deviations

| Region | Area (mm<sup>2</sup>) | | Total Voxels with Boutons | | Voxels with $\geq 30$ Boutons (Red) | | Voxels with $< 30$ Boutons (Blue) | | Total Neuronal Somata | |
| --- | --- | --- | --- | --- | --- | --- | --- | --- | --- | --- |
|  | Mean | Std Dev | Mean | Std Dev | Mean | Std Dev | Mean | Std Dev | Mean | Std Dev |
| Aud | 9.39 | 2.49 | 18.17 | 5.74 | 0.00 | 0.00 | 18.17 | 5.74 | 0.00 | 0.00 |
| CAA | 30.73 | 10.35 | 66.00 | 31.57 | 0.33 | 0.52 | 65.67 | 31.63 | 0.33 | 0.82 |
| CFAov | 1.84 | 0.24 | 111.00 | 55.81 | 17.17 | 22.61 | 93.83 | 50.59 | 3.17 | 2.32 |
| CFAr | 3.24 | 0.81 | 229.33 | 111.86 | 131.00 | 107.57 | 98.33 | 73.75 | 19.17 | 20.55 |
| FRo | 19.19 | 3.47 | 1544.67 | 212.63 | 937.17 | 306.11 | 607.50 | 355.62 | 74.17 | 43.20 |
| GZo | 13.54 | 0.79 | 278.17 | 156.17 | 13.50 | 20.60 | 265.50 | 139.57 | 3.00 | 3.52 |
| CFAc | 0.24 | 0.28 | 20.50 | 25.21 | 3.17 | 4.92 | 17.33 | 20.57 | 0.50 | 0.84 |
| IN | 14.63 | 2.17 | 530.83 | 111.06 | 107.00 | 60.27 | 423.83 | 93.60 | 2.00 | 2.00 |
| DZo | 6.16 | 1.21 | 391.33 | 84.92 | 98.50 | 60.07 | 292.83 | 60.97 | 6.17 | 3.49 |
| To | 0.69 | 0.19 | 3.50 | 3.89 | 1.00 | 2.45 | 3.50 | 3.89 | 0.00 | 0.00 |
| PIROI | 36.70 | 15.57 | 58.33 | 37.72 | 0.83 | 1.33 | 57.50 | 37.29 | 0.17 | 0.41 |
| PPCI | 3.03 | 0.82 | 66.50 | 47.67 | 1.83 | 2.48 | 64.67 | 45.65 | 0.17 | 0.41 |
| PPCm | 3.93 | 0.60 | 129.33 | 40.19 | 15.00 | 19.82 | 123.17 | 29.59 | 0.83 | 0.75 |
| PMBSF | 6.24 | 1.18 | 206.50 | 65.80 | 12.83 | 11.57 | 193.67 | 56.00 | 1.83 | 1.83 |
| PR | 2.22 | 0.98 | 154.67 | 76.78 | 49.67 | 30.53 | 103.33 | 52.01 | 2.33 | 1.75 |
| PRh | 2.77 | 1.30 | 120.67 | 93.32 | 7.67 | 8.38 | 113.00 | 87.00 | 0.00 | 0.00 |
| PV | 4.76 | 0.18 | 272.50 | 47.86 | 94.83 | 38.88 | 177.67 | 22.50 | 2.33 | 1.75 |
| RS | 1.90 | 0.69 | 21.33 | 31.41 | 1.33 | 3.27 | 20.00 | 28.50 | 0.17 | 0.41 |
| S1FPo | 1.13 | 0.43 | 49.50 | 42.56 | 3.00 | 6.00 | 46.50 | 37.04 | 0.33 | 0.52 |
| S2 | 5.83 | 0.25 | 337.50 | 58.82 | 69.17 | 39.81 | 268.33 | 26.75 | 3.50 | 4.93 |
| TP | 2.38 | 0.68 | 1.83 | 2.04 | 0.00 | 0.00 | 1.83 | 2.04 | 0.00 | 0.00 |
| Vis | 7.68 | 1.49 | 18.17 | 13.05 | 0.00 | 0.00 | 18.17 | 13.05 | 0.00 | 0.00 |

**Table 3:** Ranking of bouton densities

| Region | Score Mean | Z-scores |
| --- | --- | --- |
| FRo | 119.000 | 3.437 |
| PR | 108.833 | 2.770 |
| CFAr | 107.167 | 2.661 |
| DZ | 104.000 | 2.453 |
| CFAov | 97.667 | 2.038 |
| S2 | 97.333 | 2.016 |
| PV | 96.667 | 1.973 |
| CFAc | 81.500 | 0.978 |
| S1FPo | 78.250 | 0.765 |
| PRh | 75.000 | 0.552 |
| IN | 72.167 | 0.366 |
| PMBSF | 69.000 | 0.158 |
| PPCm | 67.500 | 0.060 |
| PPCI | 56.667 | -0.639 |
| GZ | 56.000 | -0.683 |
| RS | 32.500 | -2.224 |
| CAA | 27.000 | -2.585 |
| Vis | 26.583 | -2.612 |
| Aud | 26.167 | -2.639 |
| To | 25.083 | -2.710 |
| PIROI | 22.167 | -2.902 |
| TP | 16.750 | -3.257 |

Note: Total Voxels (with labeling) Per Region of Interest were subjected to a nonparametric test (Wilcoxon Rank Sum Test). Z-scores were used to determine cutoff values for defining regions of interest with dense, moderate and sparse connectivity. Cutoff values for z-scores were set at $\geq 2.5$ (dense), $\geq 1.0$ and $< 2.5$ (moderate), $\geq -0.5$ and $< 1.0$ (sparse). $< -0.5$ (negligible). Connectivity for regions of interest with z-scores $< 0$ were designated as negligible.

**Table 4:** Ranking of neuronal somata densities

| Region | Score Mean | Z-scores |
| --- | --- | --- |
| FRo | 123.833 | 4.025 |
| CFAr | 118.500 | 3.650 |
| CFAov | 112.167 | 3.205 |
| DZo | 103.333 | 2.584 |
| PR | 96.333 | 2.092 |
| S2 | 84.167 | 1.236 |
| GZo | 78.833 | 0.861 |
| PV | 76.500 | 0.697 |
| PMBSF | 70.833 | 0.299 |
| PPCm | 68.667 | 0.146 |
| CFAc | 64.167 | -0.158 |
| IN | 63.667 | -0.193 |
| S1FPo | 59.333 | -0.498 |
| PPCI | 43.500 | -1.611 |
| RS | 43.000 | -1.646 |
| CAA | 40.667 | -1.810 |
| PIROI | 39.833 | -1.869 |
| Vis | 39.667 | -1.881 |
| PRh | 34.000 | -2.279 |
| To | 34.000 | -2.279 |
| Aud | 34.000 | -2.279 |
| TP | 34.000 | -2.279 |

Note: Total Voxels Per Region of Interest were subjected to a nonparametric test (Wilcoxon Rank Sum Test). Z-scores were used to determine cutoff values for defining regions of interest with dense, moderate and sparse connectivity. Cutoff values for z-scores were set at $\geq 2.5$ (dense), $\geq 1.0$ and $< 2.5$ (moderate), $\geq -0.5$ and $< 1.0$ (sparse). Connectivity for regions of interest with z-scores $< 0$ were designated as negligible.

To delineate regions of interest with dense, moderate, sparse and negligible connectivity, voxel counts/mm^2^ were analyzed using a nonparametric procedure, the Wilcoxon Rank Sum Test. Significant differences in voxel counts/mm^2^ were found among the discrete regions of interest (Chi-square = 95.22, p < 0.0001). Since the number of total voxels/mm^2^ formed a rather continuous distribution, and it was not possible to delineate groups statistically, Z-scores based on the Wilcoxon test were used to derive arbitrary cut-off levels for defining densities. Regions with dense connections were defined as those with z-scores (>2.5); regions with moderate connections had Z-scores between 1 and 2.5, and regions with sparse connections had Z-scores between -0.5 and 1. Regions with z-scores < -0.5 were considered to have negligible connections.

Based on z-scores for voxel densities, FRo, CFAr, and PR had dense connectivity; DZo, CFAov, S2, and PV had moderate connectivity, while CFAc, S1FPo, PRh, IN, PPCm, and PMBSF had sparse connectivity. PPCl, GZo, RS, Vis, Aud, PirOl, and TP had negligible connectivity. Dense, moderate and sparse connections are illustrated graphically in Figure 7A.

**Figure 7.**
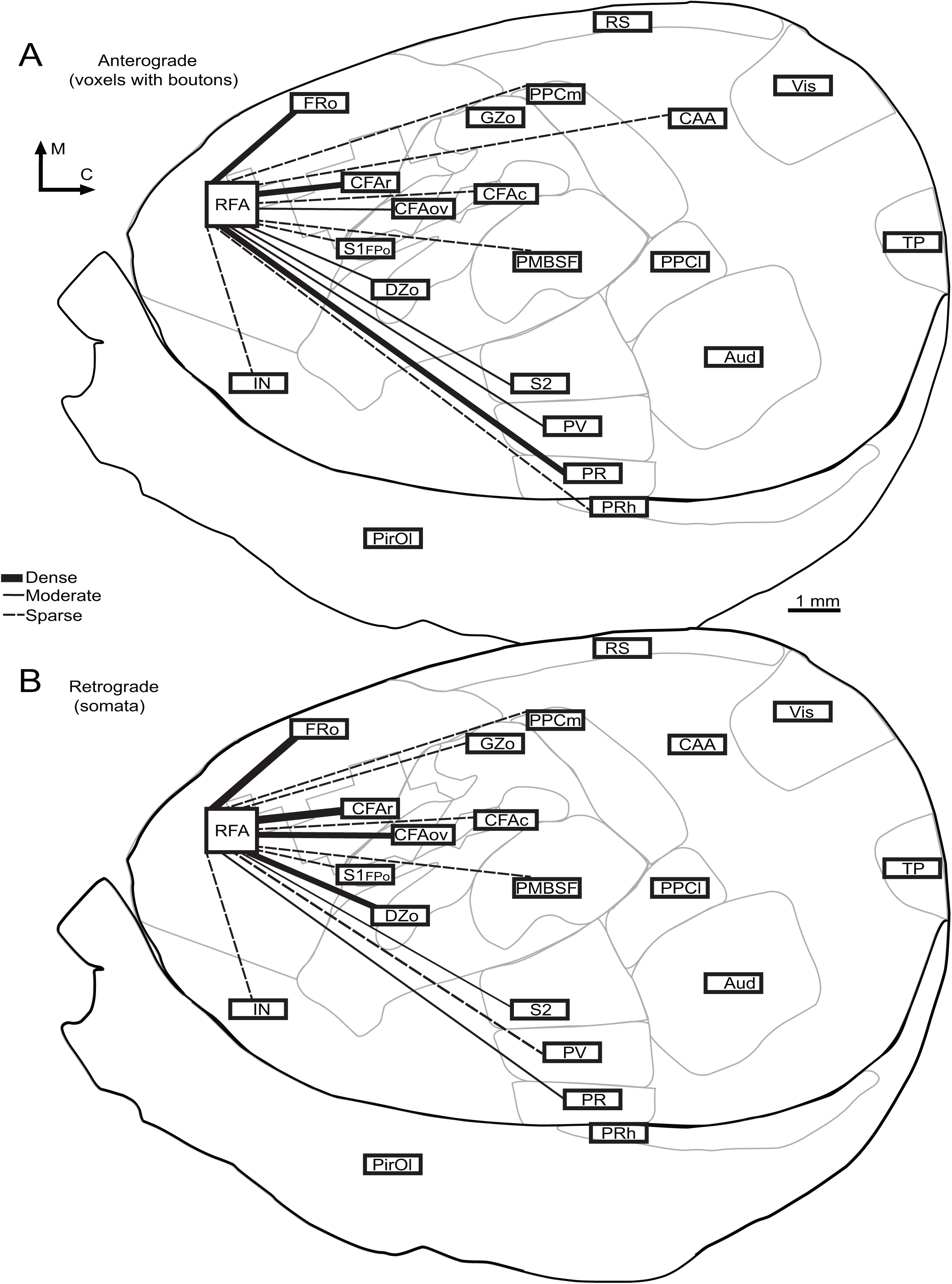
Overview of RFA corticocortical connectivity in rat. Thickness of lines indicates whether the particular connection is dense (thick black line), moderate (thin black line), or sparse (dotted line). **A**. Anterograde connectivity based on z-scores of mean voxel density across cases within each region of interest. **B**. Retrograde connectivity based on z-scores of mean somata density within each region of interest. Most areas have similar relative densities of anterograde and retrograde connections (FRo, CFAr, CFAc, PMBSF, PV, IN, S1FPo, PPCm), suggesting reciprocal connectivity; others are weighted toward retrograde connectivity (DZo, CFAov, GZo), suggesting a predominantly feed-forward connection; others are weighted toward anterograde connectivity (S2, PRh), suggesting a feedback connection.

### Quantitative Description of Labeled Somata

Total labeled somata for each region, and somata normalized to the region area, are shown in Table 2 and Figure 6, respectively. Dense, moderate, sparse and negligible connections were delineated in the same manner as voxel counts. Significant differences in somata/mm^2^ were found among the regions of interest (Chi-square = 95.22, p < 0.0001). Based on z-scores, FRo, CFAr, CFAov, and DZo had dense connectivity, PR and PV had moderate connectivity, and GZo, S2, PMBSF, PPCm, CFAc, IN and S1FPo had sparse connectivity. PPCl, RS and PirOl had negligible connectivity, while Vis, PRh, Aud and TP had no connectivity (Fig. 7B).

## Discussion

Based on tract-tracing methods describing the relative density of corticocortical termination patterns in tangential sections, the present study provides significant new details regarding the connectivity of RFA in rats. 1) RFA sends dense projections to FRo, PR and CFAr, moderate projections to DZo, CFAov, S2, and PV, and sparse projections to CFAc, S1FPo, PRh, IN, PMBSF, and PPCm (Fig. 7A). 2) As a general rule, RFA sends substantial projections to the dysgranular zone of the somatosensory cortex (DZ), but sparse to negligible projections to the granular portions of the somatosensory cortex (GZ), including the S1 forepaw area caudal to the CFA/S1 hand area overlap zone. 3) Neuronal somata projecting to RFA were dense in FRo, CFAr, CFAov, and DZo, were moderate in PR and PV, and were sparse in GZo, S2, PMBSF, PPCm, CFAc, IN and S1FPo (Fig. 7B). 3) These data may provide new insights into the functional role of RFA in motor behavior when considered in the context of the entire cortical network of sensorimotor communication.

### RFA connectivity with CFA

Similar to previous reports, RFA was found to have substantial reciprocal connections with CFA. The CFA is considered to be the forelimb representation of the rodent M1, in that forelimb movements can be evoked from this area with the lowest levels of ICMS current, and the cytoarchitecture (at least in the rostral portion) is consistent with motor cortex with respect to large layer 5 pyramidal cells. This area includes a cytoarchitectonically-distinct region coextensive with the lateral agranular field (Donoghue JP and SP Wise 1982), and called here CFAr, as well as the adjacent territory overlapping with the granular somatosensory cortex, called here CFAov due to the overlap between CFA and S1. This overlap region is also sometimes called the M1-S1 overlap zone. A large number of corticospinal neurons originate from RFA, CFAr, and CFAov. (Neafsey EJ and C Sievert 1982; Nudo RJ and RB Masterton 1990; Rouiller EM *et al*. 1993), reinforcing their role in motor behavior. In fact, a dense cluster of corticospinal neurons located in what is now called RFA was one of the original indications that a separate rostral motor representation existed in rat cortex (Hicks SP and CJ D’Amato 1977).

Previous studies have shown that the RFA and CFA are reciprocally and densely interconnected, though the laminar distribution of these connections differs substantially. Deeper layers in RFA project to CFA, while more superficial layers in CFA project to RFA (Rouiller EM *et al*. 1993). Based on these laminar connection patterns, the RFA projection to CFA has been described as a feedback connection, while the CFA projection to RFA has been described as a feedforward connection. This would appear to be consistent with a recent photostimulation study of RFA-CFA connections in mice (Hira R et al. 2013). Photostimulation of one of these areas resulted in evoked postsynaptic spikes in the other. More specifically, this photostimulation study suggests that RFA receives strong functional connections from layer 2/3 and/or layer 5a of CFA, while CFA receives strong functional connections from layer 5b of RFA.

In the present study of BDA injections into RFA, anterograde and retrograde labeling was found throughout CFA. However, the density tended to be somewhat higher in CFAr compared with CFAov. In CFAr, there was not a clear distinction in anterograde versus retrograde connectivity, as the connection was considered dense in both directions. In CFAov, however, there was some propensity for greater relative density of labeled soma compared with labeled boutons. Thus, based on relative density of retrograde and anterograde connectivity with RFA, and in line with its overlap with S1, CFAov may distinctly represent the primary feedforward component of the CFA.

While RFA and CFA are strongly interconnected, contain large numbers of corticospinal neurons and are responsive in similar ways to ICMS, suggestions have been made that they are specialized for different aspects of motor function. Reversible cooling experiments in RFA and CFA in rat have suggested that RFA is more involved in grasping and CFA in reaching (Hyland B 1998; Brown AR and GC Teskey 2014). Also, while RFA neurons have similar basal spiking properties, time-course, amplitude and direction preference during forelimb movements, RFA neurons are modulated to a greater extent based upon the behavioral context (Saiki A et al. 2014). This is consistent with the notion that RFA is at a higher level of the motor hierarchy.

### RFA Connectivity with S1

In addition to connections with CFA, the present results show that RFA has substantial connections with S1. This is in line with previous reports that demonstrated RFA terminals in all cortical layers in S1 (Rouiller EM *et al*. 1993). However, a striking result of the present study was the differential connectivity of RFA with particular subareas of S1. Connectivity with the dysgranular cortex of S1 (DZo) was moderate to dense, while connectivity with granular cortex of S1 (S1FPo, PMBSF, GZo) was sparse to negligible. This distinction was clear in each of the cases, especially with regard to terminal labeling (Fig. 5). It should be noted that in the present paper, we chose not to distinguish DZ from the transitional zone (TZ) of S1. The terminal patterns were similar in these two zones, and the collective DZ+TZ zone in squirrels has been suggested to be homologous to area 3a of primates (Cooke DF et al. 2011). For simplification, in rats, we refer simply to the dysgranular zone (DZ).

The granular and dysgranular zones of S1 differ in several respects. The most obvious difference is a reduced layer 4 in the dysgranular zone. The caudal half of the DZ receives deep proprioceptive information, while the granular zone, including S1FP, PMBSF, and GZ, receives cutaneous information (Chapin JK et al. 1987). There is also a distinction between subcortical projections that may be involved in sensorimotor integration. Terminal projections from DZ and GZ overlap within the striatum, suggesting integration of deep and cutaneous inputs. However, DZ and GZ projections are largely segregated in the thalamus (except for parts of the posterior nucleus (***should we define PO, it is not defined in table 1)), brainstem and spinal cord (Lee T and U Kim 2012).

There is one exception to the differential connectivity of S1 zones with RFA. CFAov, the portion of GZ where movements can be evoked at low current levels, had moderate (terminals) to dense (soma) connectivity with RFA. This pattern is more characteristic of DZ. Thus, RFA connectivity with M1 extends beyond the rostral portion of M1 into the overlap zone between M1 and S1. An unanswered question is whether CFAov and DZ have differential connectivity with RFA based on laminar interconnections that might support different feedforward and feedback information transfer.

### RFA Connectivity with Lateral Somatosensory Areas (S2 and PV)

In the present study, RFA displayed moderate projections to two multimodal somatosensory regions, S2 and PV, located in the lateral parietal cortex. Retrograde connectivity with these areas was sparse (S2) to moderate (PV). In the rat, both S2 and PV are somatotopically organized, and each contains a full somatosensory representation of the body. The S2 somatotopic representation is a mirror image of the representation in S1. Likewise, the PV representation is a mirror image of the S2 representation (Fabri M and H Burton 1991; Remple MS *et al*. 2003). Both S2 and PV contain neurons contributing to the corticospinal and cortico-medullary pathways (Li X-G et al. 1990). Thus, RFA is a major node processing somatosensory-motor information in higher-order somatosensory areas. In an earlier study describing corticocortical connectivity of RFA, the lateral somatosensory areas were not differentiated, but simply referred to as S2 (Rouiller EM *et al*. 1993). In that study, terminals were uniformly distributed throughout all layers in S2, similar to S1.

Given the specificity in topographic organization in S2 and PV, an unexpected finding was the seemingly non-topographic connectivity these areas have with RFA. S2 and PV are contiguous at the forelimb and hindlimb representations (Remple MS *et al*. 2003), so that a topographic pattern would be expected to be comprised of a single cluster of labeled boutons/soma located at the common forelimb representation between the two regions. Instead, in the current study, we found two distinct clusters of labeled terminals and somata lateral to S1: one in S2 and one in PV. This was most apparent for terminal labeling (Figs 1I, 5A, 5C), as retrograde labeling was more scattered and diffuse. While we did not explore the somatosensory representations in S2 and PV using neurophysiological mapping techniques, the densely connected zone appears to be located in more proximal body representations of both S2 and PV based the location of the labeled areas in their position relative to the auditory cortex (Koralek KA et al. 1990; Remple MS *et al*. 2003). This pattern was consistent across the sample of rats in the study. Further, the region where S2 and PV adjoin was largely devoid of labeled terminals, especially in the most rostral and caudal aspects corresponding to forelimb and hindlimb representations. There was one exception, illustrated in Figure 1I, where two separate clusters of densely labeled terminals were apparent near the common border of S2 and PV, presumably corresponding to the forelimb representation. This atypical connectivity pattern may suggest the importance of modulation of the proximal somatosensory representations during forelimb movement. However, clarification of the corticocortical topography between RFA and S2/PV awaits studies combining neurophysiological mapping and anterograde labeling techniques in secondary somatosensory areas.

### RFA Connectivity with frontal cortex (FR) and posterior parietal cortex (PPCm, PPCl)

FRo, as defined here, is the frontal cortex region rostral to CFA, but excluding RFA, the site of the tracer injection. That is, it includes the surrounding territory rostral, medial and lateral to RFA, as well as the motor representations intervening between RFA and CFA. Nearly half of all labeled voxels in the hemisphere were found within FRo. While there is likely to be some bias for labeled terminals and somata in the immediate surrounds of RFA simply due to the proximity to the injection site, clear projection patterns can be seen extending to territories quite medial and lateral to RFA. Extremely high projection zones were found in FRo (areas enclosed by green lines in Fig 5), especially medial to RFA and in the extreme orbitofrontal area. FR (and RFA) is contained within the medial agranular (AGm) cortex. AGm, along with the posterior parietal (PPC) cortex, is thought to be part of a cortical network for directed attention in rats (Reep RL and JV Corwin 2009). In the present study, we also found sparse connections in the medial part of the presumed PPC (PPCm), consistent with prior results (Reep RL et al. 2003). Thus, the present results confirm that RFA interacts with this network for directed attention (AGm and PPCm).

Connections of RFA with PPCl (sometimes called posterolateral or PL cortex; (Remple MS *et al*. 2003)) were considered negligible across the present sample of rats. It should be noted, though, that in some rats, a substantial number of labeled terminals were clearly observed in PPCl (Fig 5). PPCl is a multisensory region found at the caudo-lateral border of S1 barrel cortex, immediately lateral to PPCm, posterior to S2, and medial to auditory cortex (Brett-Green B et al. 2003). While precise borders were not identified using neurophysiological means in the present study, neurons in this region appear to be multisensory, and are modulated by both auditory and somatosensory stimuli (Brett-Green B *et al*. 2003; Remple MS *et al*. 2003). It is possible that PPCl represents another region with access to RFA involved in integration of multisensory information for correct stimulus-driven guidance of the forelimb.

### RFA connectivity with parietal rhinal cortex (PR)

Moderate (somata) to dense (terminals) labeling was also found in PR. Located lateral to S2 and PV, PR may relay visceral information based on its connections with brainstem and spinal cord (Cechetto DF and CB Saper 1987; Fabri M and H Burton 1991). Since this area is heavily interconnected to the S1 granular zones, it has been suggested that PR could play a role in fusion of internal and external body maps (Fabri M and H Burton 1991). The present results suggest that this network includes RFA as well.

### RFA connectivity with IN

RFA connections with IN were found to be sparse. However, it should be noted that in the present study, IN was not subdivided. The vast majority of labeled terminals and somata were located in the most rostral portion of IN, where a substantial number of voxels were found with >30 boutons/voxel (see red dots in Figure 1I, 5A, 5C, 6A). Also, localized regions with an unusually high density of terminal labeling were occasionally found in rostral IN (Figure 5A). This was the only region with such unusually high density other than FR. Thus, despite the relatively low ranking for IN as a whole, the rostral IN has substantial connections with RFA.

Rouiller et al. also reported substantial connectivity between RFA and agranular insular cortex, i.e., the most rostral portion of IN (Rouiller EM *et al*. 1993). This connection with rostral IN was one of the characteristics of RFA that distinguished it from CFA. In that study, terminal labeling in the adjacent dysgranular insular cortex was sparse, while virtually no connections were found with the more caudal granular insular cortex.

Neurons in agranular insular cortex play a role in integrating somatosensory information with chemosensory information (Katz DB et al. 2001). It has further been suggested that such chemosensory-somatosensory processing is involved in perceptual valuation of food and reward (Baldo BA et al. 2015). Taken together with the present results, there is substantial evidence that RFA has direct access to the limbic system, deriving motivational aspects of environmental cues, and integrating them with motor commands.

### RFA connectivity with PRh

In the present study, sparse terminal connections were found between RFA and PRh, though no labeled somata were found in any of the rats. These results are consistent with a previous study employing retrograde tracer injections into the perirhinal and entorhinal cortex, which reported labeled cell bodies in frontal cortical areas (Burwell RD and DG Amaral 1998). In that study, frontal cortical areas contributed about 8% of the total cortical connections to perirhinal cortex, and about half of those projections were from the so-called “secondary motor areas”, which likely includes RFA. The current study provides firm evidence that the electrophysiologically identified RFA projects to perirhinal cortex.

It was previously shown that electrical stimulation of the anterior portion of the perirhinal cortex results in evoked field potentials in frontal cortex (Kyuhou S-i and H Gemba 2002). In that same study, BDA injected into the anterior part of perirhinal cortex resulted in labeled terminals in both RFA and CFA, primarily in layers 1, 2 and 6. Since no labeled somata were found in PRh in the present study, it is possible that the terminal fields found in the Kyuhou and Gemba study were in a frontal cortical area not within the RFA.

A recent network analysis of a curated dataset derived from previous literature has provided evidence that the entorhinal cortex has the most extensive corticocortical connection pattern of all areas studied (Bota M et al. 2015). Thus, it is not surprising that such connections may exist. However, while it is well-known that perirhinal cortex, along with postrhinal and entorhinal cortex are involved in memory functions, and it can be speculated that they support motor memory, the specific role played by connections with RFA are as yet unclear.

### Similarities Between Premotor Connections in Rodents and Primates

The premotor cortex is classically defined as a region within the frontal cortex immediately anterior to M1, located within Brodmann’s area 6, and thought to represent the highest level of the motor hierarchy (Fulton JF 1935). Dum and Strick (Dum RP and PL Strick 2002) operationally defined premotor cortex as regions in the frontal lobe that project directly to motor cortex. Based on this definition, they demonstrated that in primate species, premotor cortex contains at least six (possibly up to nine) separate areas, including the supplementary motor area (SMA), the cingulate motor areas (CMA), dorsal premotor cortex (PMd) and ventral premotor cortex (PMv). Each of these premotor areas projects to the spinal cord via the corticospinal tract, and electrical stimulation of each area results in evoked movements. Thus, the RFA in rats meets these classical definitions of a premotor area. It is directly located anterior to M1 (i.e., CFA), it has dense reciprocal connections with M1, forelimb movements can be evoked with relatively low levels of ICMS current, and many of its layer 5 neurons project to the spinal cord.

From an evolutionary standpoint, there is unlikely to be a rodent homolog of premotor cortex. This would require that the areas were derived from a common ancestor. However, in a study of corticospinal neurons in two dozen mammalian species, Nudo and Masterton (Nudo RJ and RB Masterton 1990) argued that neurologically more primitive (and extant) mammals that have more distant common ancestry with primates have only two cortical areas that originate corticospinal neurons, the combined M1/S1 region and the second somatosensory region. In fact, all mammals studied had this feature in common. A separate, dense cluster of corticospinal neurons anterior to M1 was found in all primate species (in the presumed PMv), all rodent species (in the presumed RFA), but no other mammalian Orders studied (except for rabbit, in the Order Lagomorpha, typically included with the Order Rodentia in the Superorder Glires; (Douzery EJ and D Huchon 2004)). Thus, Nudo and Masterton proposed that separate premotor areas emerged independently in Primates and Glires, i.e., after the divergence of the mammalian Orders. It is possible that similar selective pressure for higher order motor control existed in the two mammalian lineages, leading to parallel or convergent evolution of premotor cortex in primates and rodents.

A persistent question in motor neuroscience is whether RFA is most similar in structure and function to any particular primate premotor area (Nudo RJ and SB Frost 2007). Alternatively, one could consider RFA an amalgam of primate motor areas, not yet evolutionarily differentiated into multiple subregions. Rouiller et al. (Rouiller EM *et al*. 1993) noted the similarities between rat RFA and monkey SMA with respect to connections with insular cortex. However, SMA in primates contains a complete topographic map of the body, while the RFA representation in rats is limited to the proximal and distal forelimb, though there are rare reports of hindlimb movements evoked from a few sites in this region (Neafsey EJ 1990).

Identical tract-tracing techniques and analytic approaches to those of the present study have been used to quantitatively define the intracortical connections of PMv in squirrel monkeys (Dancause N, S Barbay, SB Frost, EJ Plautz*, et al.* 2006). Comparing rat RFA and squirrel monkey PMv connections reveals striking similarities (See Fig 11 in (Dancause N, S Barbay, SB Frost, EJ Plautz*, et al.* 2006)). First, the cortical areas with the greatest connectivity with PMv are the rostro-lateral portion of M1 and frontal areas rostral to PMv; the connection pattern of rat RFA is similar. Cortical areas with moderate connectivity with PMv in squirrel monkey and RFA in rat include S2 and PV. Finally, PMv in monkey and RFA in rat both have sparse connections with posterior parietal cortex. Of course, there are areas connected to PMv that do not seem to have an equivalent counterpart in rats, including the additional premotor areas PMd, SMA and CMA. However, it would seem that, in general, not only do RFA and PMv have similar connection patterns qualitatively, but the order of the hierarchy with respect to the strength of the connection is nearly identical.

The parallels in connectivity between RFA and PMv with S1 deserve special attention. In squirrel monkey, PMv has moderate connectivity with area 3a, but negligible connections with area 3b. This pattern is also similar in rats, in that RFA has dense connections with the dysgranular zone of S1 (possibly equivalent to area 3a), but sparse connectivity with the granular zone of S1. Thus, despite the somewhat different arrangement of primary somatosensory areas in rodents, with granular and agranular regions more interdigitated, the primate PMv and the rodent RFA both have a connectivity preference for less granular regions.

One final similarity between the RFA and PMv is the incomplete cortical representation of body movements. RFA is surrounded by face, neck and trunk representations similar to PMv of primates. With rare exceptions, hindlimb movements cannot be evoked from RFA, PMv or the immediately surrounding cortex.

One caveat regarding the similarities between RFA and PMv connectivity is that similar quantitative data do not yet exist for PMd, SMA and CMA. Because all of the primate premotor areas are part of an interconnected network, similarities may also exist in connectivity patterns between RFA and these other primate premotor areas. Nonetheless, the present study argues strongly for RFA as a premotor area with intracortical connections strikingly similar to those of premotor areas in primates.

## Conclusion

The present study provides the strongest evidence to date that RFA is a premotor area, much like premotor cortex of primates. Its connectivity patterns with primary motor cortex, primary somatosensory cortex, secondary somatosensory areas, and posterior parietal cortex are strikingly similar. The tangential sectioning approach reveals the distinct preference for RFA connections with dysgranular zones of S1, where proprioceptive information is processed. Along with its dense connections with insular and entorhinal cortex, RFA appears to occupy a unique position in the motor hierarchy, integrating information regarding motivation and memory, as well as proprioceptive information from broad regions of the body, cutaneous information from the forelimb, and feedback from executive functions in the primary motor cortex to modulate motor behavior.

## Acknowledgments

This work was supported by the National Institutes of Health grant number R37NS030853 to RJN. The authors would like to thank Billie Byerley for her help with data collection. Corresponding author: Randolph J. Nudo, Ph.D., Landon Center on Aging, MS 1005, University of Kansas Medical Center, 3901 Rainbow Boulevard, Kansas City, KS 66160.

